# A small RNA pathway mediates global allelic dosage in endosperm

**DOI:** 10.1101/138750

**Authors:** Robert M. Erdmann, P.R. V. Satyaki, Maja Klosinska, Mary Gehring

## Abstract

Balance between maternal and paternal genomes within the triploid endosperm is necessary for normal seed development. The majority of genes in Arabidopsis endosperm are expressed in a 2:1 maternal:paternal ratio, reflecting endosperm genomic DNA content. Here we find that the 2:1 transcriptional ratio is not, unexpectedly, a passive default but is actively regulated. We describe an inverse relationship between the parent-of-origin of small RNAs and mRNAs in endosperm on a genome-wide scale. Disruption of the Pol IV small RNA pathway causes the entire transcriptome to become more maternally biased. Furthermore, paternal inheritance of a RNA Pol IV mutation is sufficient to rescue seed abortion caused by excess paternal genome dosage. These results indicate that maintenance of the maternal:paternal transcriptome ratio in endosperm is an active process and reveal a function for RNA Pol IV in mediating the global transcriptional balance between maternally and paternally inherited genomes in endosperm.

## Introduction

Proper endosperm development is required for the formation of viable seeds. Seeds of flowering plants consist of three genetically and epigenetically distinct components: the two products of fertilization, diploid embryo and triploid endosperm, and diploid seed coat derived from maternal ovule tissues. Whereas the embryo has a typical 1:1 ratio of maternal to paternal genomes, the ratio of maternal to paternal genomes in the endosperm is 2:1. Most genes expressed in the endosperm mirror the 2:1 maternal:paternal genomic DNA ratio, with the exception of a subset of 100-200 genes that are imprinted and are expressed primarily from maternal or paternal alleles (Gehring et al., 2011; Hsieh et al.,2011). DNA methylation and chromatin modifiers have previously been shown to regulate the maternal to paternal transcript ratios of imprinted genes (Hsieh et al., 2011). Active DNA demethylation occurs in the central cell, the female gamete that is fertilized to produce the endosperm, before fertilization (Park et al., 2016). Consequently, after fertilization maternally inherited genomes are hypomethylated compared to paternally inherited genomes, particularly at transposable elements and their remnants (Gehring et al., 2009; Hsieh et al, 2009). Differential DNA methylation between maternal and paternal alleles is one important feature of many imprinted genes in multiple plant species (Gehring and Satyaki, 2017). Additionally, there is strong evidence that trimethylation of H3K27 by PRC2 (Polycomb Repressive Complex 2) maintains imprinted expression patterns in endosperm (Gehring et al., 2006; Jullien et al., 2006; Hsieh et al., 2011; Wolff et al., 2011; Zhang et al., 2014; Moreno-Romero et al., 2016). Variations on these features also exist (Klosinska et al., 2016). It is unknown whether or not there are regulators of the maternal to paternal transcript ratios at the majority of genes that are expressed in endosperm but not imprinted.

Small RNAs have diverse functions in eukaryotes, including post-transcriptional gene silencing and transcriptional gene silencing *via* guided modifications to DNA or chromatin (Borges and Martienssen, 2015). DNA methylation in all sequence contexts (CG, CHG, and CHH) is established and maintained by the RNA-directed DNA methylation pathway. The most abundant class of plant small RNAs are 24 nucleotide (nt) small RNAs (sRNAs), which primarily emerge from transposable elements and other repetitive genomic regions (Matzke and Mosher, 2014). In the canonical RNA-directed DNA methylation (RdDM) pathway, the biogenesis of 24 nt small RNAs is initiated by RNA Polymerase IV (Pol IV). Pol IV produces short non-coding transcripts that are converted into double-stranded RNA by the interacting RNA-dependent RNA polymerase RDR2, and are then diced by DCL3 into 24 nt small RNAs. 24 nt small RNAs can be loaded into AGO proteins to guide the DRM2 *de novo* DNA methyltransferase to matching DNA sequences, establishing transcriptional gene silencing by RdDM (Matzke and Mosher, 2014). Several variations to the canonical pathway exist, and 24 nt small RNAs can also arise independent of Pol IV transcription (Cuerda-Gil and Slotkin, 2016).

Small RNAs are highly abundant during the reproductive phase of the *Arabidopsis thaliana* life cycle (Mosher et al., 2009). Previous studies have reported genome-wide, parent-of-origin specific small RNA profiles from Arabidopsis seed pods (Mosher et al., 2009) or whole seeds (Pignatta et al., 2014). Imprinted genes that are maternally expressed in the endosperm are associated with paternally biased 24 nt small RNAs in whole seeds (Pignatta et al., 2014) and wild-type paternal *NRPD1* in somatic tissues is required for silencing of the paternal allele for two maternally expressed imprinted genes (MEGs) in endosperm (Vu et al., 2013). Expression of the gene encoding the largest subunit of RNA Pol IV, *NRPD1*, is paternally biased in endosperm at 6 days after pollination (DAP) in *A. thaliana*, with on average 55% (range of 43-69%) of transcripts from the paternally inherited allele (compared to the expectation of 33%) (Gehring et al., 2011; Pignatta et al., 2014). In the related species *A. lyrata*, expression of gene encoding a common subunit of Pol IV and Pol V, *NRPD4*, is paternally biased (Klosinska et al., 2016). These findings suggest that the paternally inherited genome promotes RdDM, perhaps to counteract consequences of maternal genome hypomethylation. Despite the possible connections to imprinting, the precise relationship between small RNAs in Arabidopsis endosperm and maternal or paternal allele expression or methylation remains unknown. Here we investigated the distribution, parent-of-origin, and function of small RNAs in *Arabidopsis thaliana* and *Arabidopsis lyrata* endosperm. These experiments uncovered genome-wide control of maternal:paternal transcriptional ratios, mediated by small RNAs, at the majority of endosperm-expressed genes.

## Results

### Increased accumulation of genic small RNAs in endosperm

To determine the identity and distribution of small RNAs in the two products of fertilization, we performed small RNA sequencing from multiple, replicated wild-type genotypes of *A. thaliana* embryos (n=14 independent libraries) and endosperm (n=16 independent libraries) isolated at 6 days after pollination (**Table S1**). This generated around 250 million mapped sRNA reads for each tissue. 24 nt sRNAs were the most abundant size class in both embryo and endosperm, although endosperm had a somewhat higher fraction of 21 nt small RNAs (**Fig. S1**). Most embryo and endosperm sRNAs mapped to pericentric chromosome regions (Fig. 1a), as has been previously described for other Arabidopsis tissues (Greaves et al., 2012). An additional sRNA population associated with the more gene-rich chromosome arms was observed in endosperm (Fig.1a). Genome-wide, transposable element sequences were associated with relatively lower levels of sRNAs in endosperm compared to embryos, and gene bodies were associated with relatively higher levels (Fig. 1b). We confirmed that the unique endosperm small RNA profile was not due to contamination from the maternal seed coat (**Fig. S2**) and was dependent on RNA Pol IV (Ye et al., 2016) (**Fig. S3**).

**Figure 1:**
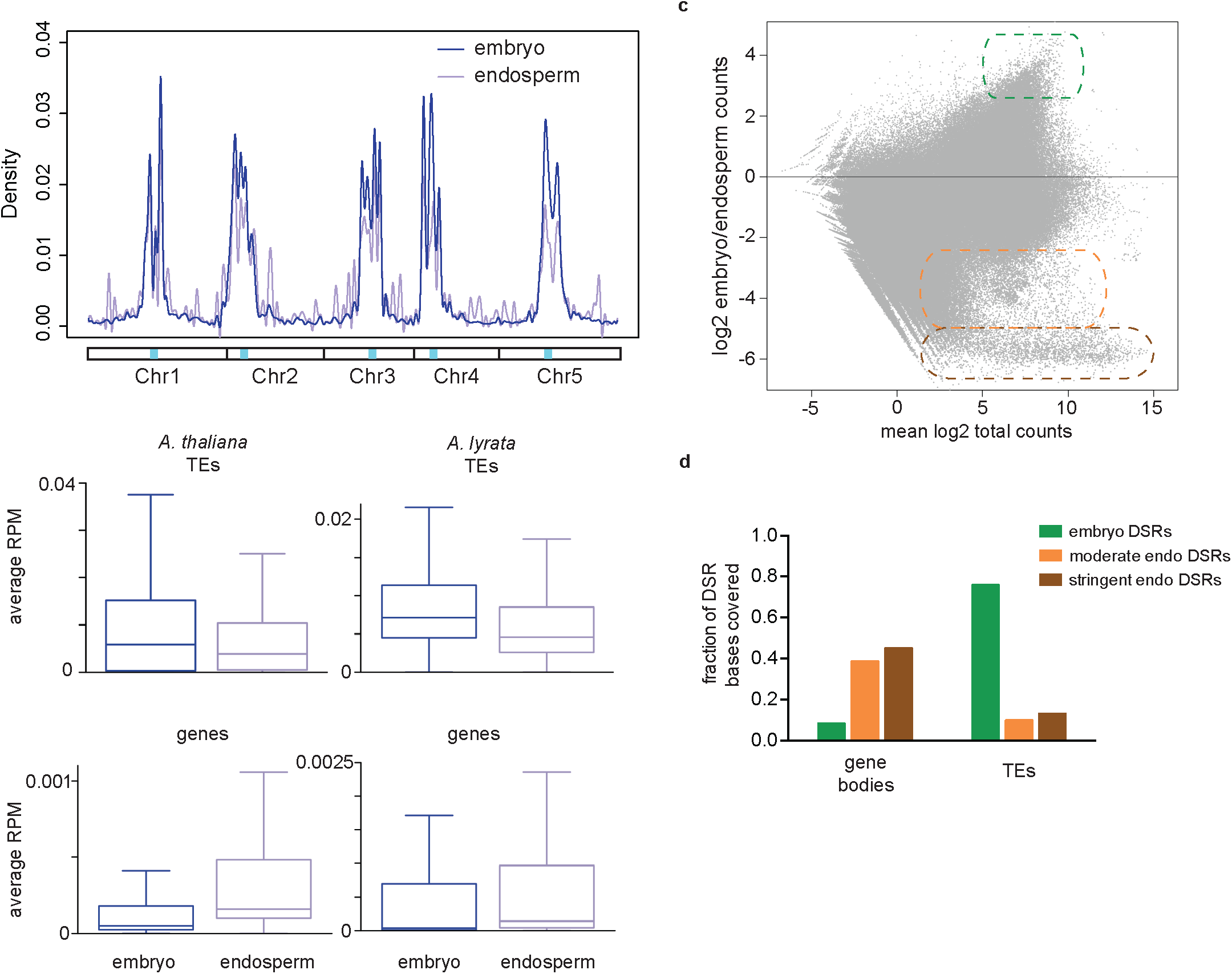
Endosperm has a unique small RNA profile. **a**, Kernel density plot of *A. thaliana* embryo and endosperm 24 nt small RNA abundance genome-wide. Blue shading within chromosomal schematic below the plot shows the relative position of the centromeres. **b**, Box and whisker plots of the average RPM of 24 nt sRNAs in TEs (top) and gene bodies (bottom) for *A. thaliana* (left) and *A. lyrata* (right) embryo and endosperm. Line is the median; whiskers are 1.5x the interquartile range. **c**, MA plot illustrating the identification of three categories of differential sRNA regions (DSRs) for 21-24 nt *A. thaliana* small RNAs. Small RNA values were determined for overlapping 300 bp windows throughout the genome. Green outline, embryo DSRs; orange outline, moderate endosperm DSRs; brown outline, stringent endosperm DSRs. **d**, Overlap of DSRs with gene bodies and TEs. See also Figures S1, S2, and S3.

To determine the relevance of our findings outside of *A. thaliana*, we also profiled small RNAs in embryo (3 replicates) and endosperm (3 replicates) of the outcrossing perennial *A. lyrata*, resulting in about 120 million mapped reads per tissue. Similar features were observed. Small RNAs corresponding to annotated TEs were relatively more abundant in embryo compared to endosperm, and small RNA corresponding to genes were relatively more abundant in endosperm (Fig. 1b).

To identify regions that showed tissue-enriched sRNA expression, termed here differential sRNA regions or DSRs, we binned the 21-24 nucleotide fraction of the *A. thaliana* sRNA data into overlapping 300 bp windows (Fig. 1c). DSRs were at least 2.5-fold more enriched in one tissue, and consisted of three types: embryo DSRs (n=335), moderately-enriched (“moderate”) endosperm DSRs (n=3640) or stringently-enriched (“stringent”) endosperm DSRs (n=818) (Fig. 1c, see Experimental Procedures for definitions). The genomic distribution of embryo and endosperm DSRs was distinct from one another and consistent with genome-wide trends (Fig. 1b): embryo DSRs were most prevalent in transposable elements and endosperm DSRs were more prevalent in genes (Fig. 1d).

### Endosperm-enriched 24 nt small RNAs are not consistently associated with DNA methylation

In other tissues, 24 nt small RNAs are associated with DNA methylation of cognate sequences, a hallmark of which is the accumulation of non-CG methylation. We determined whether regions with differential small RNA accumulation were differentially methylated between embryo and endosperm (Fig. 2). Consistent with expectations, in *A. thaliana* regions corresponding to embryo DSRs were more methylated in the embryo than endosperm in all cytosine contexts. By contrast, stringent and moderate endosperm DSRs were associated with no difference in CG methylation and with only minor increases in endosperm non-CG DNA methylation (Fig. 2a,c). These results indicate that A. thaliana endosperm-enriched small RNAs do not always correlate with increased DNA methylation. *A. lyrata* displayed a different relationship between 24 nt small RNAs and methylation. We previously showed that like *A. thaliana*, *A. lyrata* endosperm is CG hypomethylated on maternal alleles compared to the embryo. However, unlike *A. thaliana*, there are also many regions of maternal allele CHG hypermethylation, which are associated with reduced maternal allele expression (Klosinska et al, 2016). We found that like *A. thaliana*, regions less CG methylated in *A. lyrata* endosperm than embryo were associated with lower small RNA abundance in endosperm (Fig. 2b). Regions that were CHG hypermethylated in endosperm compared to embryo were associated with higher levels of endosperm small RNAs. These results suggest that 24 nt small RNAs in *A. thaliana* and *A. lyrata* endosperm may operate through both DNA methylation dependent and independent pathways.

**Figure 2:**
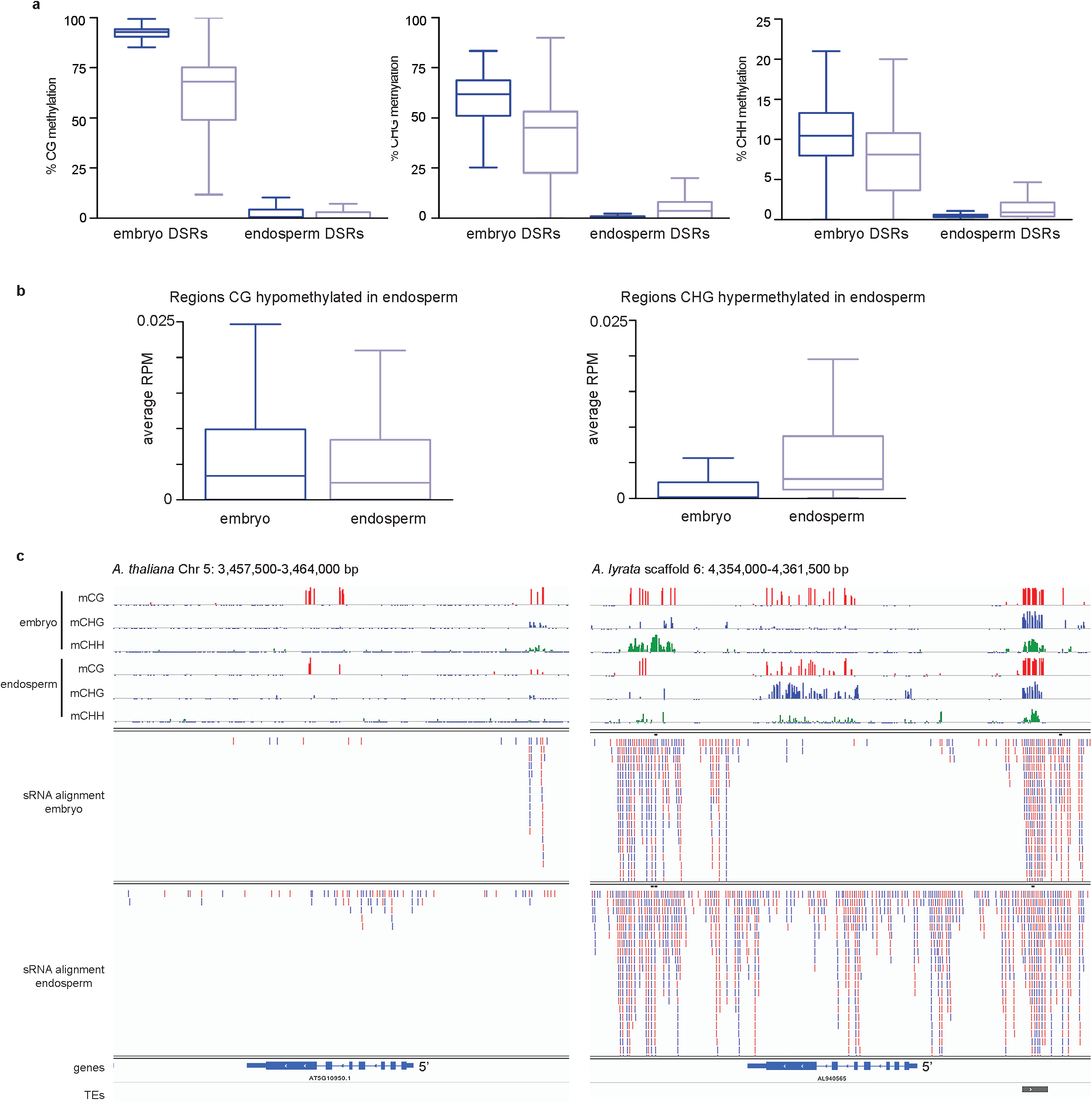
Relationship between small RNAs and DNA methylation. **a**, % CG, CHG, and CHH methylation in A. thaliana embryo DSRs and stringent endosperm DSRs in embryo (dark purple) and endosperm (light purple) 6 days after pollination. **b**, sRNA abundance in *A. lyrata* regions with less CG methylation in the endosperm than embryo and at regions with greater CHG methylation in endosperm than embryo. **c**, *A. thaliana* L*er* × L*er* embryo and endosperm small RNA (21-24 nt) and DNA methylation profiles at the PEG AT5G10950 (left) and at the syntenic region in *A. lyrata* (right) from Kar × MN crosses. Methylation scales from 0-100%. Tickmarks below the line indicate a cytosine with bisulfite sequencing coverage but no methylation. *A. thaliana* methylation data from Pignatta et al., eLife 3: e03198 (2014); *A. lyrata* methylation data from Klosinska et al., Nat Plants. 2: 16145 (2016). Note that *A. thaliana* and *A. lyrata* sRNA alignments are not scaled to one another. For *A. thaliana* approximately 14 million reads are aligned for each tissue. For *A. lyrata* approximately 60 million reads are aligned for each tissue.

### Endosperm small RNAs are globally biparentally expressed, but biased towards maternal allele expression in genes

Several of the profiled small RNA populations were derived from a cross between two wild-type polymorphic parents (**Table S1**), allowing examination of the contribution of maternally and paternally inherited genomes to small RNA biogenesis. The genome-wide proportion of maternal and paternal small RNAs was similar to the respective ratios of maternal:paternal genomes in each tissue: 60% of endosperm small RNAs and 50% of embryo RNAs were derived from the maternally inherited genome in *A. thaliana* (Fig.3a). Similarly, 42% of embryo small RNAs were derived from maternal alleles in *A. lyrata*, as were 54% of endosperm small RNAs. These data are consistent with small RNA data from Arabidopsis whole seeds (Pignatta et al., 2014) and from rice and maize endosperm (Rodrigues et al., 2013; Xin et al., 2014), in which both maternally and paternally derived small RNAs have been described. Maternally and paternally derived embryo small RNAs exhibited identical genomic distributions (Fig. 3b) and very few parentally biased embryonic small RNA regions were identified (**Fig. S4**). The vast majority of regions that produced endosperm small RNAs were not parentally biased (Fig. 3c). However, about 10% of regions were strongly biased towards expression from the maternally or paternally inherited genome (Fig. 3c). Regions that accumulated greater than 90% paternal small RNAs in endosperm (2.96% of all regions that could be evaluated) tended to be found in traditionally heterochromatic regions of the genome and were not differentially expressed between embryo and endosperm (Fig. 3d). Maternally derived endosperm small RNAs were found both in heterochromatic regions and in gene-rich chromosome arms (Fig. 3b) but the majority of regions associated with greater than 90% maternal small RNAs in endosperm (6.54% of all regions that could be evaluated) were expressed more highly in the endosperm than in embryos (Fig. 3d) and included stringent endosperm DSRs (Fig. 3e). Maternally derived small RNAs were enriched at the 5′ and 3′ ends of the transcribed unit of genes (Fig. 3f). These data suggested that there might be a relationship between small RNAs and allelic expression in endosperm.

**Figure 3:**
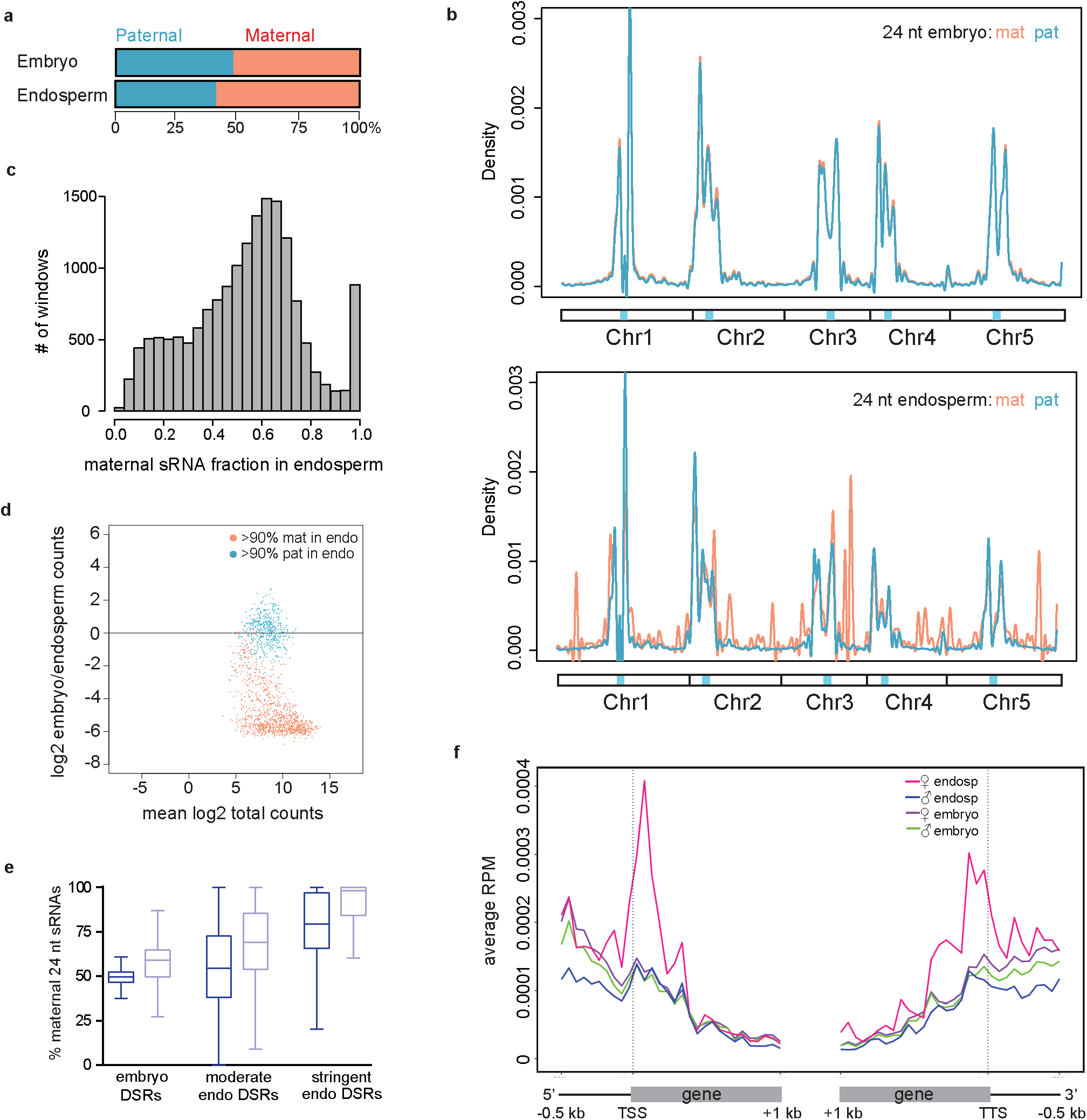
Endosperm sRNAs are produced from both maternally and paternally inherited genomes, with some regions of parental bias. **a**, Average % sRNAs derived from maternally and paternally inerhited genomes in embryo (n=8 libraries) and endosperm (n=10 libraries). **b**, Kernel density plot of maternally and paternally derived 24 nt sRNAs in embryo and endosperm. **c**, Histogram of endosperm 24 nt maternal sRNA fraction for 16,747 300 bp regions with sufficient data. **d**, MA plot illustrating the expression patterns of endosperm sRNA regions that are over 90% maternal (red) or paternal (blue) in origin. **e**, Box and whiskers plots of the % maternal sRNAs in DSRs in embryo (dark purple) and endosperm (light purple). Line represents the median. Whiskers are 1.5x the interquartile range. **f**, Metaplot of maternally and paternally derived 24 nt sRNAs over the 5’ and 3’ portion of genes and flanking regions in embryo and endosperm. All data from *A. thaliana*. See also Figure S4.

### Parental small RNA bias is inversely correlated with parental mRNA bias

To determine if there was a general relationship between the maternal:paternal ratios of endosperm small RNAs and mRNAs, we examined the maternal fraction of mRNAs per gene based on the maternal fraction of overlapping 24 nt small RNAs (Fig.4, **Fig. S5**). We previously reported that maternally expressed imprinted genes were enriched for paternally-derived small RNAs in whole seeds (Pignatta et al., 2014). Consistent with this, we found that maternally expressed imprinted genes were associated with paternally biased endosperm small RNAs in the gene body and flanking regions (**Fig. S5**). Conversely, paternally expressed imprinted genes were associated with maternally biased small RNAs, particularly in the gene body and 3’ flanking regions. 80% of paternally expressed imprinted genes overlapped a stringent endosperm DSR. Expanding the analysis to all genes revealed a more general negative association between maternal sRNA fraction and maternal mRNA fraction. The maternal mRNA fraction for genes decreased as the maternal sRNA percentage associated with those genes increased (Fig. 4a). These data suggest that there is a reciprocal relationship between small RNA and mRNA production that is not restricted to imprinted genes, but occurs throughout the genome.

**Figure 4:**
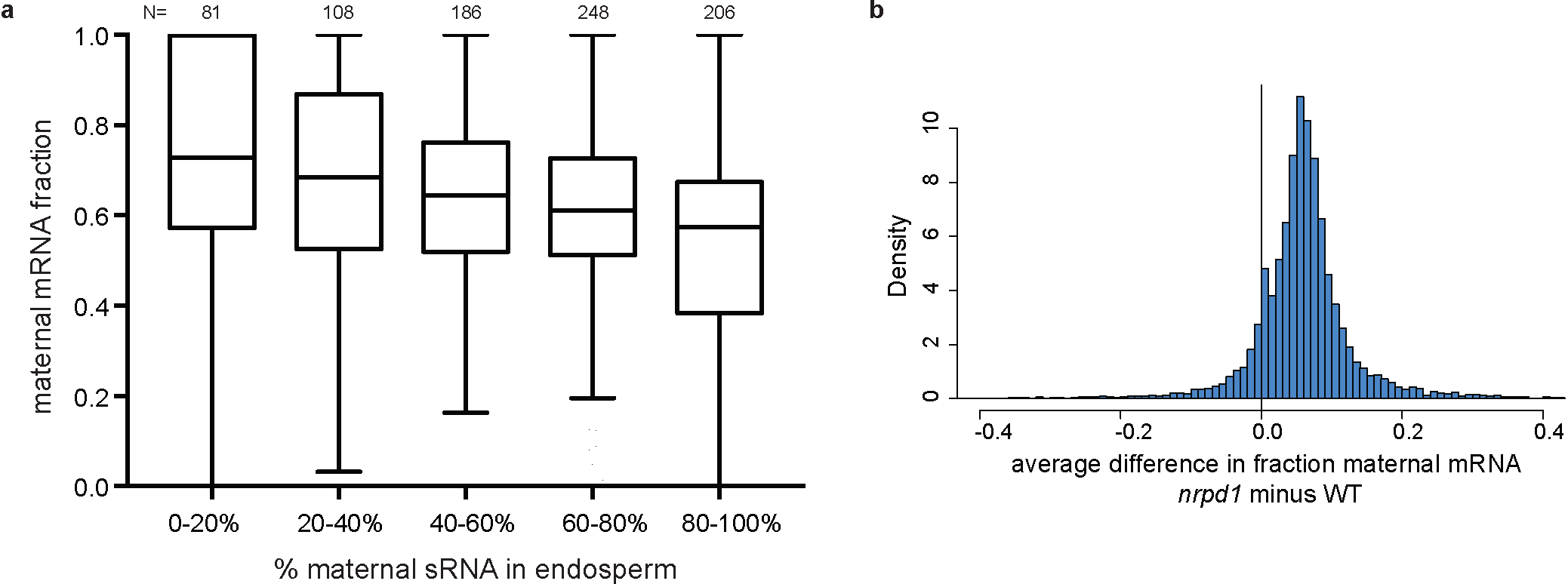
Loss of Pol IV is associated with a genome-wide shift towards maternal allele mRNA expression in endosperm. **a**, Maternal sRNAs are negatively associated with maternal mRNAs. Box and whisker plots show the maternal fraction of mRNAs per gene in each sRNA category for wild-type Ler × Col crosses. Number of genes in each sRNA category indicated above plot. Line represents the median. Whiskers are 1.5x the interquartile range. **b**, Histogram of the difference between *nrpd1* and wild-type (WT) maternal mRNA fraction for 11,003 individual genes in endosperm. All data from *A. thaliana*. See also Figure S5.

### The ratio of maternal:paternal transcripts is shifted towards the maternal allele in RNA Pol IV mutants

If the negative association between the parent-of-origin of 24 nt small RNAs and mRNAs is causal, eliminating Pol IV-dependent sRNAs should affect the relative ratio of maternal to paternal allele transcripts. *NRPD1* encodes the largest subunit of RNA Pol IV (Onodera et al., 2005). We examined parent-of-origin specific gene expression in *nrpd1* mutant endosperm by mRNA-seq (**Table S2**). The expression of many paternally expressed imprinted genes shifted towards the maternal allele in *nrpd1* mutants, consistent with repression of the maternal allele by *cis*-produced small RNAs in wild-type (**Fig. S5**). However, we found that this phenomenon was not specific to imprinted genes. Rather, in *nrpd1* mutant endosperm the expression of most genes shifted toward the maternal allele by on average around 5-10% compared to the wild type (Fig. 4b, **Fig. S5**). These data were confirmed by pyrosequencing cDNA of 9 different loci from wild-type and mutant crosses. For 8 of 9 genes examined from multiple independent replicates, the average fraction of transcripts from the maternal allele was higher in *nrpd1* mutant endosperm compared to the wild type (**Table S3**). These results indicate that mutations in Pol IV result in a global shift in transcriptional balance towards maternally-inherited alleles.

### Mutations in RNA Pol IV suppress paternal genomic excess phenotypes

To explore the consequences of the transcriptional shift towards the maternal genome, we altered the relative contribution of maternal and paternal genomes in wild-type and *nrpd1* mutant endosperm. Seeds with a 1:1 ratio of maternal:paternal genomes in endosperm often abort (Lin et al., 1984; Scott et al., 1998; Pennington et al., 2008). Given the excess of global maternal allele expression in *nrpd1* mutant endosperm, we reasoned that *nrpd1* mutants might suppress the phenotypic effects of increased paternal genomes. Tetraploid *nrpd1* mutants were created by colchicine treatment and confirmed by FACS analysis. Seed abortion and seed germination phenotypes from crosses between wild-type or *nrpd1* mutant parents were assessed (Fig. 5). As expected, crosses between wild-type diploid females and wild-type tetraploid males (producing endosperm with a 1:1 maternal:paternal ratio) resulted in 81% aborted seeds, with another 10% exhibiting an abnormal phenotype (Fig. 5a,b). By contrast, when diploid *nrpd1* females were crossed to tetraploid *nrpd1* males, seed abortion was reduced to 24% (Fig. 5a,b), indicating substantial rescue. 85% of tested seeds from this cross germinated (Fig 5c) and were confirmed to be triploid. We also investigated whether having one parent mutant for *NRPD1* was sufficient to suppress the seed abortion phenotype. Seeds lacking only the paternal *NRPD1* allele, which is more highly expressed than the maternal allele in diploid endosperm, also displayed reduced seed abortion, to 13%, and were able to germinate and form true leaves (Fig. 5). By contrast, inheritance of the *nrpd1* allele from the diploid female parent only did not suppress seed abortion in a cross to a wild-type tetraploid male (Fig. 5b,c). Together, these data suggest that the negative effects of an extra paternal endosperm genome might be suppressed by the relatively higher maternal allele expression caused by mutations in *NRPD1*, although seed rescue may be dependent on expression differences of individual genes.

**Figure 5:**
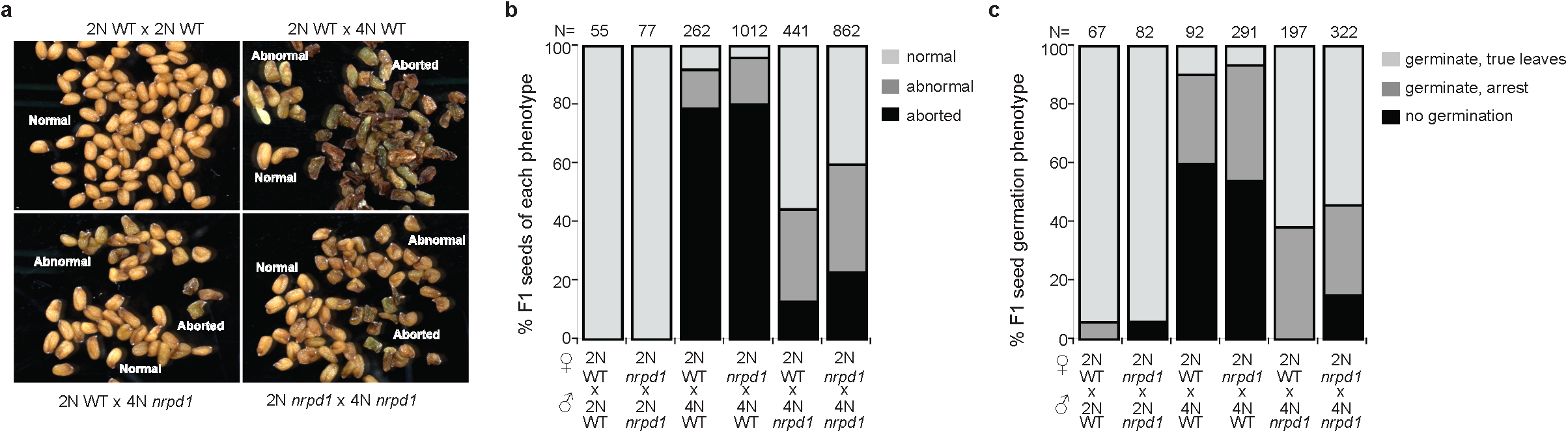
Mutations in NRPD1 suppress paternal genomic excess phenotypes. **a**, Seed phenotypes observed in crosses beween A. thaliana wild-type (WT) diploids, WT tetraploids, and diploid and tetraploid *nrpd1* mutants. Female in the cross listed first. Scale bar is 500 µm. **b**, Quantification of F_1_ seed phenotypes observed in the indicated crosses. **c**, Seed germination of F_1_ seeds from the indicated crosses. Seeds that germinated but did not develop beyond the cotyledon stage were classified as arrested. Number (N) of seeds analyzed for each cross indicated above the plot for b and c.

### Discussion

Transcriptional dosage control has been described with regard to the ratio of sex chromosomes to autosomes and in polyploids and anueploids (Disteche, 2016). The application of allele-specific mRNA-seq to endosperm several years ago (Gehring et al.,2011; Hsieh et al., 2011) suggested that the endosperm was not subject to parent-of-origin specific global dosage control – most genes were expressed in a ratio reflective of genomic DNA content. This study has revealed that maintaining the expected 2:1 ratio is an active process mediated by RNA Pol IV. In its absence, expression shifts towards more expression from maternally inherited genomes. Thus our results have revealed previously hidden transcriptional dosage control. Several questions remain unanswered, foremost among them why an active process is required to maintain a seemingly default transcriptional ratio. One possibility is that this is a necessary response to the alterations of the maternal epigenome that are initiated before fertilization. Maternally-inherited endosperm DNA is hypomethylated due to active DNA demethylation before fertilization, an essential process that establishes gene imprinting (Huh et al., 2008; Gehring and Satyaki, 2017). Additionally, endosperm chromatin is decondensed compared to chromatin from other cell types, and there is some evidence that this is specific to the maternally inherited genomes (Baroux et al., 2007; Pillot et al., 2010). The combined effect of DNA demethylation and chromatin decondensation could render the maternally inherited genomes more permissive to transcription, and thus in the absence of an active mechanism expression from maternal genomes would be favored. Our findings raise the possibility that hidden dosage control may also exist in other systems, particularly when genomes or chromosomes are in distinct epigenetic states.

The precise mechanism by which 24 nt small RNAs exert their effects on genes in the endosperm remain unknown. Regions of the genome with the highest levels of small RNAs in endosperm, like TEs, correspond to the most highly methylated regions of the genome (Fig. 1a), suggesting that RdDM is functional in endosperm. By contrast, although genic endosperm-enriched small RNAs are associated with DNA hypermethylation in *A. lyrata*, the methylation gains in *A. thaliana* endosperm are quite modest (Fig. 2). Thus while DNA methylation is often an indicator of Pol IV activity, our A. thaliana data suggests that the transcriptional effects we observed may not be dependent on DNA methylation.

The overall significance of our findings is highlighted by the ability of mutations in a Pol IV subunit to suppress seed abortion caused by excess paternal genomes (Fig. 5) – the first developmental phenotype ascribed to *nrpd1* mutants. Our findings place *NRPD1* among a relatively small number of genes that when mutated can paternally suppress triploid seed abortion (Kradolfer et al., 2013; Schatlowski et al., 2014; Wolff et al., 2015; Huang et al., 2017). Although the precise molecular basis for seed abortion suppression by *nrpd1* remains unknown, our data suggest that a critical function of Pol IV is during reproductive development.

Parental conflict is thought to underlie the evolution of gene imprinting in flowering plants and mammals (Haig, 2013). The idea behind this theory is that maternally and paternally inherited genomes have differing interests with regard to maternal investment in offspring (in plants, endosperm), the outcome of which is parent-of-origin specific (imprinted) gene expression. Our data suggest that there are two outcomes of conflict in endosperm, operating at different scales: the first is at individual imprinted genes and the second at the level of the entire genome. RdDM pathway genes, which themselves are imprinted, function at both of these scales. The phenomenon described here is likely to be broadly relevant – endosperm from interspecific hybrid tomato seed have phenotypes suggestive of maternal:paternal genomic imbalance and display a global shift towards expression from maternally inherited genomes (Florez-Rueda et al., 2016). In conclusion, our results demonstrate that the Pol IV sRNA pathway mediates dosage interactions between maternal and paternal genomes, highlighting an active tug-of-war between parents with regard to transcriptional outcomes in the endosperm.

## Experimental Procedures

### Plant material

*A. thaliana* plants were grown in a greenhouse with 16-hr days at approximately 21°C. For crosses, flowers were emasculated two days prior to pollination. To introgress the *nrpd1a-4* allele into the L*er* background, we performed a series of 4 backcrosses of the SALK_083051 line from Col into the L*er* background followed by self-fertilization to acquire homozygous *nrpd1a-4* mutants. Col and *nrpd1a-4* tetraploids were generated by the application of 0.25% colchicine in 0.2% Silwet to the apices of 2-3 week-old diploid plants. Tetraploid progeny of these plants were identified by flow cytometry. To test germination for Fig. 5C, seeds were sterilized and plated on 0.5x MS/Phytagar media. The seeds were monitored for germination for up to two weeks after plating. *A. lyrata* plants of the Kar and MN47 strains (Klosinska et al, 2016) were grown in a growth chamber with 16-hr days at 20°C.

### Seed dissection and RNA isolation

*A. thaliana* seed dissections occurred 6 days after pollination, generally corresponding to the torpedo stage of embryogenesis under our growth conditions. *A. lyrata* seed dissections occurred at the walking stick stage of seed development at 15 days after pollination. RNA was isolated from manually dissected endosperm and embryo, as described (Gehring et al., 2011), using the RNAqueous Micro Kit (Ambion, Life Technologies Corporation, Carlsbad, CA). The small RNA pool was separated from the larger RNA pool by sequential ethanol addition and filtering. The larger RNA pool was collected on filters after the addition of ethanol equivalent to 60% of the sample volume, and the small RNA pool was collected on filters after the addition of ethanol equivalent to 50% of the flow-through volume from the larger RNA filter-binding step.

### Small RNA library construction

Libraries for Illumina sequencing were constructed using the NEXTflex Small RNA-Seq Kit v2 as directed (Bioo Scientific Corporation, Austin, Texas). 40 base single-end sequencing of sRNA libraries was performed on an Illumina HiSeq machine (**Table S1**).

### Small RNA analysis

Analysis of sRNA libraries began with trimming low-quality read ends (fastq_quality_trimmer, -t 20 and -l 25; http://hannonlab.cshl.edu/fastx_toolkit/). Adapters were removed with cutadapt (−0 6 -m 26 --discard-untrimmed) (Martin, 2011). sRNA libraries prepared using the NEXTflex sRNA-Seq Kit v2 are appended with 4 base randomized barcodes immediately 3’ and 5’ of the sRNA read itself. These barcodes were used to remove PCR duplicates (any read of the same sRNA sequence with identical flanking barcodes on both ends), before being removed themselves. Following adapter and barcode removal, the resulting reads were analyzed with fastqc (http://hannonlab.cshl.edu/fastx_toolkit/).

Reads were aligned using Bowtie 1.1.1 (Langmead et al., 2009) with two mismatches allowed (-v 2 –best), using a metagenomic approach to reduce mapping bias. In the case of crosses between Col and L*er* ecotypes, a metagenome was constructed using the TAIR10 genome and the published L*er* genome (Gan et al., 2011). In the case of crosses between Col and Cvi ecotypes, a metagenome was constructed using a Cvi pseudogenome (Cvi 1bp indels and SNP substitutions introduced into the TAIR10 genome) and the TAIR10 genome. Following alignment, reads aligning to either parent genome were converted back to TAIR10 coordinates, and were classified by strain using SNPs (Pignatta et al., 2014). If classification at two SNP positions within the same read conflicted, that read was discarded. All reads that overlapped annotated tRNAs, snRNAs, rRNAs, or snoRNAs were removed. For *A. lyrata*, reads were aligned to the MN47 reference genome (Hu et al., 2011).

With a final set of aligned reads, a single base read depth value for sRNAs of any desired size class was created – each size of sRNA was normalized separately so that each base of a read n-nucleotides in length contributed 1/n of a read to an overlapping genomic position. Alignment files were run through bedtools genomecov (-d -scale: adjusted to account for individual library size; bedtools v. 2.23.0, bedtools.readthedocs.io/en/latest/index.html). Per-position values for all libraries pooled within a given analysis were averaged, and the resulting per position average values could then be intersected with any genomic locus of interest using bedtools intersect (bedtools v. 2.23.0). This single base averaging method was used for all non-MA plot-derived analyses, which instead utilized a windows-based approach described below.

### DSR calling

In order to identify DSRs, *A. thaliana* 21-24 nt sRNAs were binned into 300 bp windows with 150 bp overlaps, such that exactly 2 windows covered every position in the genome. Overlapping was performed using bedtools coverage (bedtools v. 2.23.0).
sRNA window read values, classified as either embryo or endosperm, were used as input for DESeq2, with the sizeFactors command used to provide RPM normalization between libraries (Love et al., 2014). The 14 embryo and 16 endosperm individual library output values were then used to calculate mean sRNA levels for all libraries at a given locus (mean log_2_ total counts) and the ratio of mean embryo expression to mean endosperm expression (log_2_ embryo/endosperm counts), and resulting values were used as the basis for MA plot construction. Embryo DSRs were defined as higher expression of 21-24 nt sRNAs in embryo libraries than in endosperm libraries, with an FDR-adjusted p-value of <1×10^−10^, and a log_2_ embryo/endosperm counts value of greater than 2.5. Moderate endosperm DSRs have the same p-value requirement as embryo DSRs, only with a log_2_ embryo/endosperm counts value of less than −2.5 and greater than −5. Stringent endosperm DSRs have the same p-value requirement, but require a log_2_ embryo/endosperm counts value of less than −5.

### mRNA library construction

2 µg of DNase I-treated RNA was used to prepare mRNA-seq libraries using the Truseq Stranded mRNA Library Prep Kit as directed (Illumina Inc., San Diego, CA). Amplification was for 15 cycles. Libraries were sequenced with 100×100 base paired end reads on an Illumina HiSeq machine (**Table S2**).

### mRNA data analysis

Analysis of mRNA reads began with read filtering, using cutadapt as described for sRNAs, and trim_galore (-t 10; http://www.bioinformatics.babraham.ac.uk/projects/trim_galore/) to remove low quality bases from the 5’ end of reads. Trimmed reads were assessed with fastqc. Reads were aligned to TAIR10 using Tophat v2.0.13 (-N 2 -read-edit-dist 2) (Kim et al., 2013). Following alignment, reads were classified by strain using SNPs as described above for sRNAs. The same general procedure was followed for both single-end and paired-end reads.

In the case of libraries originating from crosses including a parent with the *nrpd1a-4* allele in the L*er* background, reads originating from genes that retained the Col genotype even after multiple backcrosses into L*er* were removed before downstream analysis. To accomplish this, reads that fell within loci that had a log_2_ ratio of Col to L*er* allele calls above 2.5 in both directions of a reciprocal cross were excluded. This was done separately for each of the three replicates, which were derived from different backcross progeny. The primary regions of Chromosome 1 that were excluded from analysis included AT1G48410 through AT1G80990 for replicate 1, AT1G48410 through AT1G70420 for replicate 2, or AT1G48410 through AT1G77720 for replicate 3.

### Pyrosequencing

Categories of loci tested with pyrosequencing included highly expressed genes identified as maternally shifted within the *nrpd1* mRNA-seq data (AT3G45010, AT2G17980), imprinted genes that are maternally expressed in wild-type (AT2G17690, AT5G38140, AT4G31060), imprinted genes that are paternally expressed in wild-type (AT3G45090, AT5G11460), and loci identified as paternally shifted within the *nrpd1* mRNA-seq data (AT1G08600, AT3G05400). Col/L*er* SNPs selected for testing within these loci were not flanked by other SNPs for at least 100 bp in either direction. PCR amplification of loci was performed using the Qiagen PyroMark PCR kit as directed. The University of Michigan DNA Sequencing Core performed pyrosequencing reactions. Individual assays where the non-SNP-specific nucleotide signal was greater than 10% were excluded from further analysis (marked as N/A in **Table S3**).

## Data Availability

All sequencing data will be available in NCBI GEO under record GSE94792.

## Supplemental Information

Supplemental information includes 5 supplemental figures and three supplemental tables.

## Author Contributions

RME, PRS, and MG conceived and designed experiments. RME performed the majority of experiments and data analysis, PRS performed interploidy expreiments, and MK performed *A. lyrata* experiments. All authors analyzed data. RME and MG wrote the paper with contributions from all authors.

## Acknowledgments

We thank Colette L. Picard and Prat Thiru for advice and assistance with bioinformatics analyses. This research was funded by NSF grant 1453459 to M.G. R.M.E. was supported by an NSF Graduate Research Fellowship.

